# *SbPIP2* Mediated Improvements in Plant Resilience: Physiological and Molecular Insights into Abiotic Stress Response

**DOI:** 10.1101/2024.02.13.580036

**Authors:** Jaykumar Patel, Kusum Khatri, Nirmala Kumari Gupta, Jalak Maniar, Deepesh Khandwal, Babita Choudhary, Dylan Wyn Phillips, Huw Dylan Jones, Avinash Mishra

**Author notes:** Corresponding author; (AM).

## Abstract

Understanding the mechanisms behind plant resilience to abiotic stresses is essential for enhancing crop yield and sustainability. This study integrates findings from a comprehensive investigation into the function of the *SbPIP2* gene, which encodes an aquaporin protein, in improving the abiotic stress tolerance of transgenic plants. Our integrated approach revealed that transgenic plants overexpressing *SbPIP2* significantly reduce reactive oxygen species (ROS) accumulation and exhibit enhanced physiological attributes, including higher seed germination rates, improved growth, early flowering, and better seed setting under stress conditions. Notably, these plants also showed a quicker recovery and completion of their lifecycle post-stress treatment. The transcriptomic analysis provided a deeper understanding of the genetic modifications contributing to stress resilience, highlighting the involvement of genes associated with oxidative stress response, calcium and sugar signaling pathways, stomatal regulation, phytohormone biosynthesis, and flower development. Additionally, the study underscores the central role of abscisic acid (ABA) in mediating stress responses through hormonal regulation, with transgenic plants displaying increased ABA levels due to the upregulation of biosynthesis genes and downregulation of catabolism genes. This hormonal adjustment is critical for stomatal closure, reducing water loss, and enhancing tolerance to abiotic stresses. Our findings elucidate the complex genetic and molecular pathways that underpin abiotic stress tolerance in plants, offering valuable insights for future research aimed at improving crop resilience through genetic engineering, thereby addressing the challenges of climate change and environmental stressors.

## Introduction

Climate change poses a significant threat to global agriculture, leading to erratic rainfall, extreme heat waves, and increased atmospheric CO_2_ levels. These changes contribute to various abiotic stresses like drought, salinity, extreme temperatures, and nutrient deficiencies, severely affecting plant growth and productivity (Raza et al., 2020; Atkinson et al., 2012). Plants, unable to move away from these stressors, face challenges such as water scarcity and salt contamination, which are detrimental to crop production. It’s crucial to enhance water-use efficiency and salt tolerance in crops through breeding and biotechnological approaches. Abiotic stresses can cause up to US$170 billion year^−1^ loss, emphasising the need to identify and understand crops resilient to these stresses (Razzaq et al., 2021). Studying the biochemical responses of crops to abiotic stress is key to uncovering plant resistance mechanisms (Abdelrahman et al., 2015). This research is vital for developing strategies to mitigate stress impacts, such as creating stress-resistant crops, ensuring food security and sustainable agriculture in the face of climate change (Costa et al., 2019).

In the study of plant water relations, both osmotic adjustment and the control of water permeability at cellular and tissue levels are crucial for maintaining water potential gradients and the intensity of water movement (Maurel et al., 2021). Extensive research has demonstrated the dynamic nature of plant water transport and its molecular constituents in response to a wide range of environmental and hormonal stimuli over various timescales (Aroca et al., 2012). Aquaporins, predominantly located in the plasma and intracellular membranes of most plant cells, are pivotal in these mechanisms. They facilitate water movement between cells and, to a lesser extent, manage osmotic regulation within individual cells (Maurel et al., 2015). Aquaporins have increasingly been recognised as key targets within the membrane, influenced by environmental and hormonal signals that affect plant water dynamics.

Recent studies have elucidated the role of aquaporins, particularly AtPIP2;1 in Arabidopsis, in the ABA-induced stomatal closure and osmotic water permeability in guard cells. This is evident from studies using epidermal peels of Arabidopsis, where AtPIP2;1 was found essential for ABA-induced but not light- or CO_2_-induced stomatal movements (Grondin et al., 2015). Additionally, AtPIP2;1 is crucial for the ABA-triggered increase in osmotic water permeability in guard cell protoplasts and the accumulation of hydrogen peroxide (H_2_O_2_) within guard cells (Rodrigues et al., 2017). This research highlights the dual function of aquaporins in water and H_2_O_2_ transport, which is crucial for stomatal regulation. Additionally, SnRK2.6/OST1 kinase’s involvement in ABA signaling through AtPIP2;1 phosphorylation and NADPH oxidase activation has been discovered (Grondin et al., 2015, Maurel et al., 2016). However, contrasting ABA responses in different plant parts suggest complex regulatory mechanisms (Ceciliato et al., 2019), indicating the need for further investigation into aquaporin functions under stress conditions.

Halophytes exhibit unique mechanisms for abiotic stress tolerance and are sources of stress-responsive genes (Mishra and Tanna, 2017). *Salicornia brachiata*, an edible halophyte indigenous to India’s coastal marshes, has been extensively studied for its innate ability to withstand high salinity. Research employing genomic, metabolomic, ionomic, and lipidomic approaches has led to the discovery and characterisation of novel genes pivotal in ion and redox balance, nuclear signaling, and osmoprotection (Patel et al., 2022; Mishra et., 2015; Tiwari et al., 2014; Mishra et al., 2013; Jha et al., 2009). Notably, recent investigations (unpublished) have shown that overexpressing the *SbPIP2* aquaporin gene from *S. brachiata* enhances photosynthetic efficiency and abiotic stress resilience in transgenic plants, primarily through altering physical and biochemical parameters. However, the precise mechanisms by which *SbPIP2* contributes to stress tolerance remain unclear. In this study, we employ transcriptomics, both targeted and untargeted metabolomics, in vivo reactive oxygen species (ROS) measurement, and plant physiological analyses to unravel the molecular networks influenced by *SbPIP2* gene overexpression in transgenic tobacco.

## Materials and Methods

### Plant Material

*Nicotiana tabacum* (cv. Petit Havana) seeds as a wildtype (WT), Tobacco plants transformed with empty pCAMBIA1301 as a vector control (VC), and previously characterised transgenic lines overexpressing *SbPIP2* gene (unpublished data) were used for his study. Mature S. brachiata Roxb. seeds were harvested from the seaside of Bhavnagar, Gujarat (21°73.5262"N; 72°25.2581"E) and used for *SbPIP2* gene isolation.

### Seed germination

Seeds from control (WT and VC) and transgenic tobacco lines underwent surface sterilisation via a sequential treatment of 70% ethanol and 2% NaOCl solutions, followed by five rinses using sterile water. Each petri dish featuring Murashige and Skoog (MS) basal media and MS basal media supplemented with NaCl or mannitol, was distributed into seven sections. Precisely 20 seeds from both control and transgenic lines were allocated to each section and grown for 30 days. Germination counts were meticulously conducted every other day, and the gathered data was employed to determine various seed germination metrics, including germination energy, germination percentage, and relative salt injury. Upon completion of the 30-day interval, photographic documentation of all Petri dishes was performed to establish a visual archive of the experimental results.

### Growth and Recovery of Control and Transgenic Plants under Soil-Based Stress Conditions

Seeds of control (WT and VC) and transgenic plants were surface sterilised and cultivated for 20 days on MS basal plates. Seedlings were then transferred to an autoclaved soil mixture using forceps, watered, covered with a polyethene bag, and maintained in a culture room environment. After 7-8 days of acclimatisation, the polyethene bag was removed, plants were watered every other day, and allowed to grow for an additional 20 days. In total, 40-day-old plants (20 days on plates and 20 days in soil) were subjected to stress treatments.

One set of controls (WT and VC) and transgenic plants remained untreated as a control. For salinity treatment, a 200 mM NaCl solution (in water) was used for watering every other day, while for drought stress, watering was halted. Abiotic stresses were applied for 25 days, with photographs taken on the 25th day of stress treatment. Following the stress treatment, plants were allowed to recover through regular watering every other day. Photos were documented on the 10th and 70th days of recovery.

### Plant growth analysis

Control and transgenic seeds were subjected to germination under selective antibiotic conditions. Seedlings aged 25 days were then grown in a hydroponic culture system, encased within a plastic bag to facilitate acclimatisation. The hydroponic system was maintained with a half-strength Murashige and Skoog (MS) medium, which was refreshed every five days. Growth conditions included a 16-hour photoperiod at 100 μM photons m^−2^ s^−1^, an 8-hour dark phase, a consistent temperature of 24 ± 1°C, and a relative humidity of 60%. After a period of 7-8 days, the plastic bags were incrementally removed, allowing the plants to complete their life cycle within the hydroponic culture system. Plant development was documented, and heights were measured at 4 to 6 days intervals following the 45-day mark to monitor growth rates. Upon completion of the life cycle, the number of seed pods per plant was quantified.

### Abiotic stress treatments in hydroponics culture

Sixty-day-old plants (25 days on MS plates and 35 days in the hydroponic culture system) were exposed to abiotic stresses. Salinity stress was induced by supplementing half-strength MS media with NaCl to achieve final concentrations of 100 mM NaCl + MSB and 200 mM NaCl + MSB. Drought stress was induced by adding mannitol to half-strength MS media to yield final concentrations of 150 mM mannitol + MSB and 300 mM mannitol + MSB. Plants were subjected to stress for 24 hours. Post-stress treatment, the third and fourth leaves from the shoot tip were promptly collected, immersed in liquid nitrogen, and stored at -80 °C for subsequent experimentation.

### *In-vivo* localization of hydrogen peroxide (H_2_O_2_) and superoxide radicals (O_2_ ^•−^)

The accumulation of H_2_O_2_ and O_2_^•−^ radicals under stressed and unstressed conditions was assessed in both control (WT and VC) and transgenic plants following Patel et al., 2022. Leaf discs (15 mm dimeter) from treated and control were incubated in an NBT solution (0.8 mM NBT in 50 mM phosphate buffer, pH 7.8) for 6 hours in the dark to detect O2^•−^ radicals. For in vivo localisation of H2O2, leaf discs (15 mm diameter) were immersed in a DAB solution (1 mg mL^−1^ DAB, 0.1% Triton X-100, 10 mM Na2HPO4, pH 3.8) and incubated for 6 hours in the dark. After this incubation, the leaf discs were subjected to light for another 2 hours. Subsequently, the leaf discs were bleached with ethanol to enable visualisation of the staining spots. The intensity of the staining spot were estimated using ImageJ software. In this process, staining in the outer 2 mm diameter due to tissue damage during cutting was cropped and not considered for staining spot intensity calculation.

### H_2_O_2_ and lipid peroxidation

A total of 500 mg of leaf tissue from stressed and unstressed plants [control (WT and VC) and transgenic plants] was harvested, homogenised in liquid nitrogen, and subsequently incubated in a solution containing 0.25 mL of 0.1% trichloroacetic acid (TCA), 0.5 mL of 1 M KI, and 0.25 mL of 1 M phosphate buffer at pH 8.0. The samples were maintained at 4°C for 10 minutes. Following incubation, the reaction mixtures were centrifuged at 12,000 rpm for 15 minutes at 4°C. The absorbance was then measured at 350 nm and compared against a standard curve of H_2_O_2_.

For lipid peroxidation assessment, 200 mg of harvested leaf samples were extracted using 0.1% TCA solution. Malondialdehyde content was estimated through derivatisation with 0.65% thiobarbituric acid in 20% trichloroacetic acid, along with a control reaction utilising 20% trichloroacetic acid to eliminate nonspecific measurements of anthocyanins following Jha et al., 2021.

### Microarray analysis

Total RNA was extracted from 100 mg leaf samples of stressed and unstressed plants [three replicates of each control (WT) and transgenic (L1) plants] using the RNeasy Plant Mini kit (Qiagen, Germany). Sequentially first strand cDNA, second strand of cDNA and cRNA were synthesised using Affymetrix kits (Affymetrix, USA). Single-stranded cDNA was synthesised from purified cRNA, after which the cDNA was fragmented, labelled, and hybridised with a whole-gene tobacco array according to the manufacturer’s user manual (Affymetrix, USA). Arrays were then washed and stained using the FS450_0001 protocol on a Fluidics station preloaded with stain cocktails 1 and 2 and an array holding buffer. The prepared cartridges were scanned using an Affymetrix chip scanner and the GeneChip command console program. Data were exported in CEL and Excel file formats for subsequent gene expression analysis following Patel et al., 2021.

### Untargeted metabolite profiling by GC-MS analysis

Leaf tissues (100 mg) from stressed and unstressed (WT and transgenic L1, L2, L3) plants were homogenised in liquid nitrogen, mixed with ice-cold methanol, and heated at 70°C for 10 minutes and sonication for 10 minutes. Post-centrifugation at 12,000 rpm for 10 minutes, the supernatant was treated with chloroform to eliminate pigments, then dried in a vacuum concentrator. In the derivatisation phase, the dried extract was dissolved in methoxyamine hydrochloride (60 µL, 20 mg/mL in pyridine), incubated at 37°C for 2 hours, followed by MSTFA addition and further incubation. Ribitol (6 µg) was the internal standard. The derivatised samples underwent GC-MS analysis using a Shimadzu QP2010. A 1 µL sample was injected into an FID/capillary column with helium as the carrier gas in an RTX5 MS column. Temperatures started at 80°C for 2 minutes and escalated to 315°C over 40 minutes. Peak identification was based on the NIST mass spectra library, enabling qualitative and quantitative analysis with the internal standard.

### Statistical analysis

The number of independent experiments and replicates were mentioned in the experimental procedure or figure legends as required. Most statistical analysis was performed in IBM SPSS 27 using one-way ANOVA (Tukeys HSD test) or student T-test. Most graphs were plotted in Graphpad Prism 9. Untargeted metabolite analysis was performed with Metaboanalyst 5.0 and XTSTAT premium by Lumivero.

## Results

### Overexpression of *SbPIP2* accelerates seed germination and plant growth under both unstressed and stressed conditions

Germination percentage (GP), germination energy (GE), and relative stress injury (RSI) were assessed for control (WT and VC) and transgenic plant seeds under both unstressed and stress conditions (salinity and drought). Under unstressed conditions, the GP was found to be similar for both control and transgenic lines (Figure 1A). However, when subjected to salinity and drought stress, control seeds exhibited a marked reduction in GP (38-68% for salinity stress and 44-69% for drought stress) compared to their unstressed counterparts. In contrast, transgenic lines displayed only a 3-5% reduction in germination under these stress conditions. The GE results paralleled those of the GP, with no significant differences observed between control and transgenic lines under unstressed conditions (Figure 1B). However, under stress conditions, both groups showed a decrease in GE, with control plants experiencing a more substantial reduction compared to transgenic lines under both salinity and drought stress. Notably, transgenic lines demonstrated a significantly smaller reduction in GE under drought stress, particularly at 150 mM mannitol, where their GE was comparable to that observed under unstressed conditions. As anticipated, no RSI was detected in either control or transgenic plants under unstressed conditions (Figure 1C). In contrast, under salinity (100 mM NaCl and 200 mM NaCl) and drought (150 mM mannitol and 300 mM mannitol) stress conditions, control plants exhibited a 4-8 fold increase in RSI, whereas transgenic lines showed significantly lower RSI levels.

**Figure 1:**
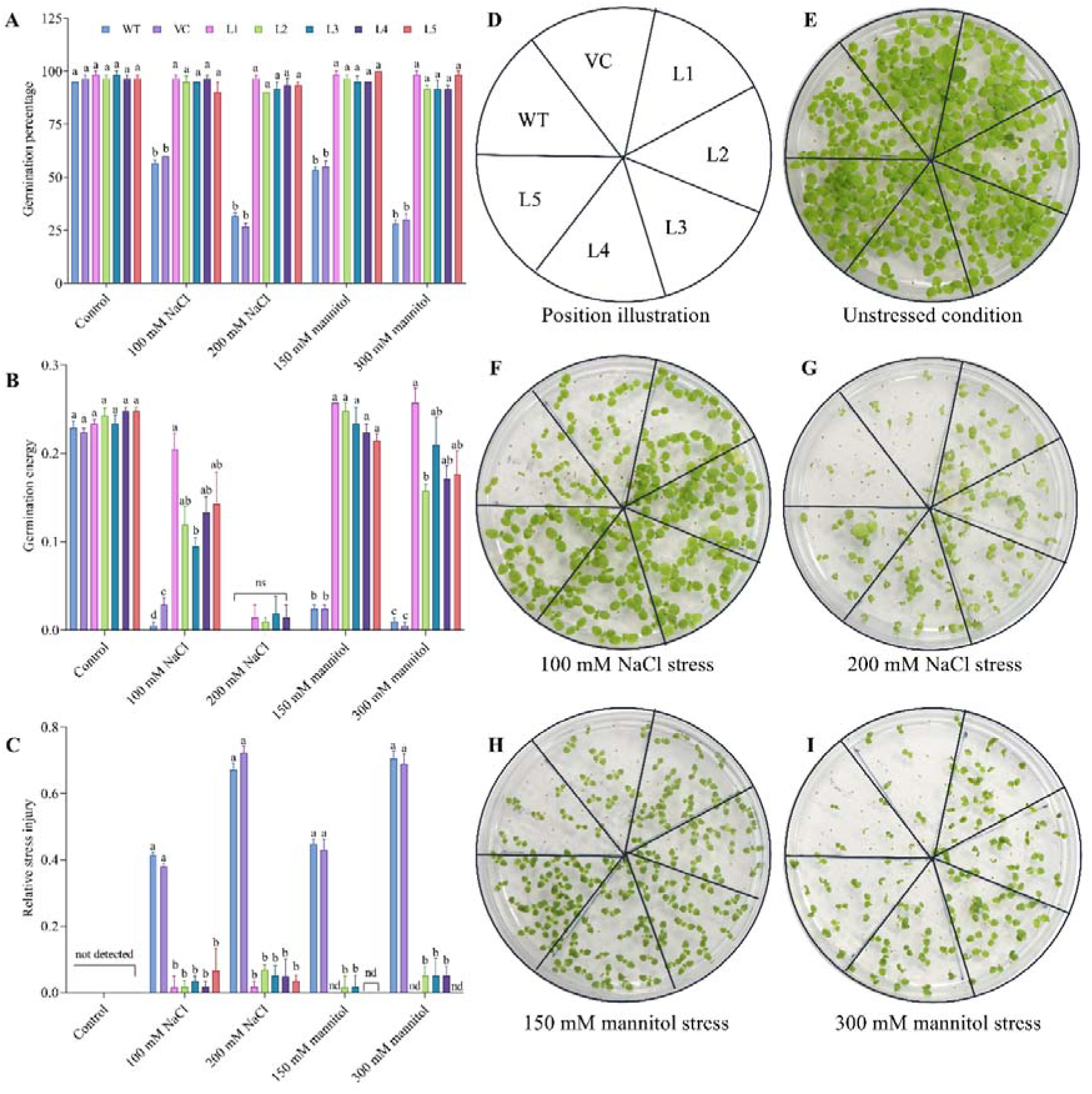
Seed germination efficiency under various conditions: This figure presents the seed germination efficiency of plants subjected to unstressed conditions, as well as different levels of salinity and drought stress, observed over a 30-day period. It includes: (A) A graph depicting the percentage of germination. (B) A graph illustrating germination energy. (C) A graph showing the relative stress injury. (D) A schematic showing the arrangement of seeds in plate. (E) Seed germination on MS basal medium, representing the unstressed condition. (F) Germination on MS basal medium supplemented with 100 mM NaCl. (G) Germination on MS basal medium supplemented with 200 mM NaCl. (H) Germination on MS basal medium supplemented with 150 mM Mannitol. (I) Germination on MS basal medium supplemented with 300 mM Mannitol. Data represent mean values (±SE) from three independent experiments, each with at least three biological replicates (n≥9). Different lowercase letters indicate statistically significant differences as determined by ANOVA, followed by Tukey’s HSD test (p < 0.05). The abbreviations used are as follows: WT - wild type, VC - vector control, L1 to L5 - transgenic lines L1 to L5, ns - not significant, nd - not detected.

Seed germination parameters were further supported by photographs documented after 30 days on MS media with or without NaCl and mannitol supplementation (Figure 1D-I). These images clearly illustrated the differences in seed germination between control and transgenic lines under salinity and drought stress. No significant differences in seed germination were observed between the groups on control MS basal plates. However, under severe abiotic stress conditions (200 mM NaCl and 300 mM mannitol), only a few control seeds germinated, while transgenic lines exhibited more successful germination and growth, albeit with altered seedling morphology due to stress. Even under milder stress conditions (100 mM NaCl and 150 mM mannitol), control seeds germinated more slowly and exhibited hampered growth compared to transgenic lines.

In this study, the growth rates of transgenic and control plants (WT and VC) were compared by cultivating them in hydroponic culture media under optimal conditions throughout their entire life cycle (Figure S1A-B). At the flowering stage, no significant differences were observed in the total length between control and transgenic plants. However, transgenic plants exhibited a more rapid growth rate, reaching their maximum height within 70 days, in contrast to control plants, which required 90-95 days. Additionally, transgenic plants demonstrated earlier flowering, with an average onset at 60 days post-germination, compared to the control plants, which initiated flowering after 90-100 days. Consequently, transgenic plants displayed accelerated maturation, seed settling, and an overall reduction in their life cycle by 30 to 40 days compared to control plants. Furthermore, a substantial increase in the number of seed pods per plant was observed in transgenic plants, with an average of 12-15 pods per plant (Figure S1C). In comparison, WT and VC plants produced only 4-6 seed pods per plant. These findings highlight the significant differences in growth rates, flowering times, and seed pod production between transgenic and control plants.

### SbPIP2 overexpression improved plant physiology in soil under both unstressed and stressed condition

Figure showed compelling evidence that salinity stress exerts a substantial negative impact on the growth of both control (WT and VC) and transgenic plants, with the latter group exhibiting a marginally lesser degree of susceptibility (Figure 2). A parallel trend was observed under drought stress conditions, where the growth of control (WT and VC) plants was notably impeded in comparison to their transgenic counterparts.

**Figure 2:**
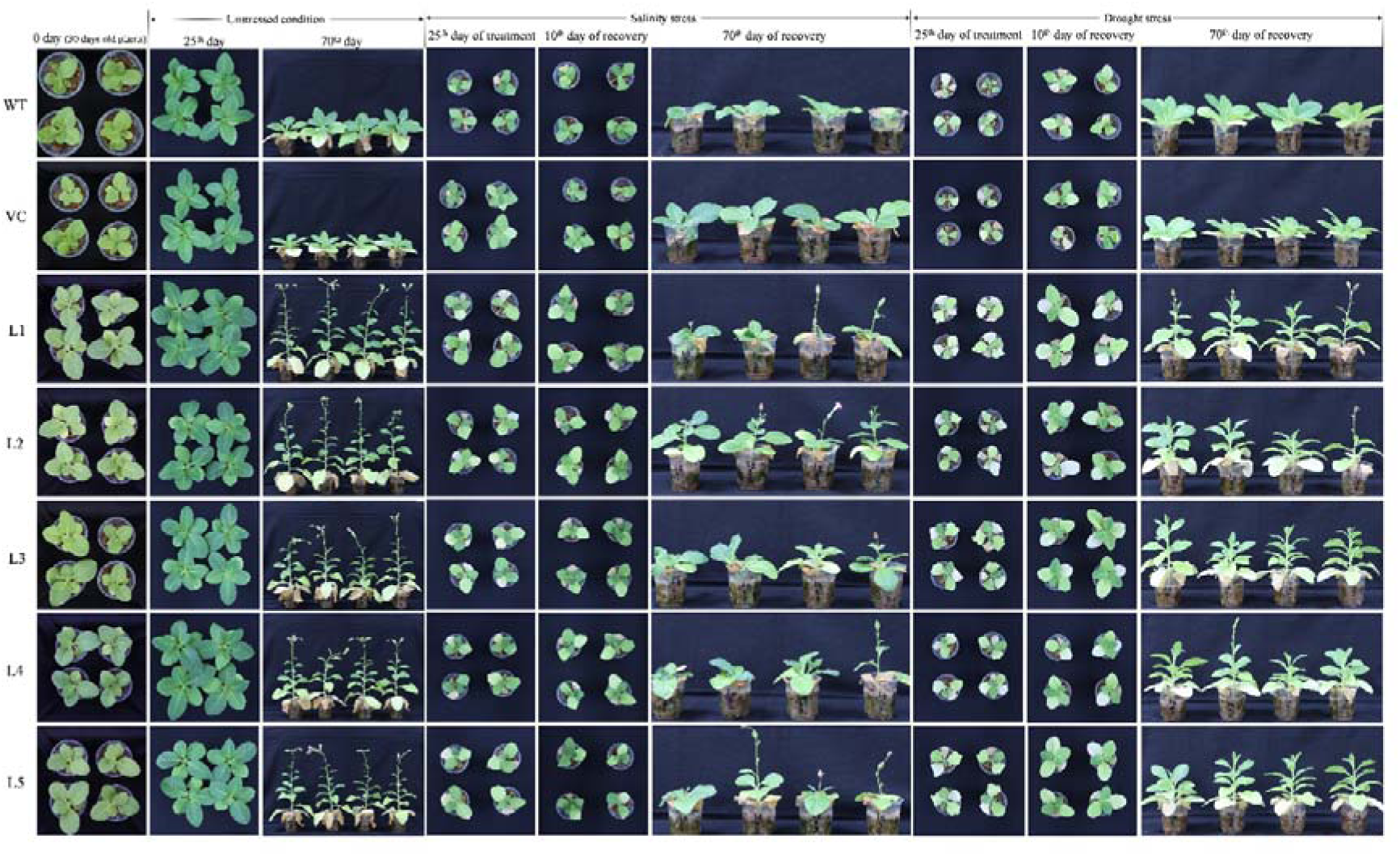
Physiology of control and transgenic plants in soil: This figure illustrates the physiological responses of 30-day-old control and transgenic plants subjected to unstressed conditions, salinity stress (200 mM NaCl solution), and drought stress (withholding water) for 25 days. Recovery was assessed by resuming regular watering (every other day) after the stress period and documenting at two intervals: 10 days and 70 days post-recovery. For plants under unstressed conditions, regular watering was maintained every other day, with documentation occurring on the 25th and an additional observation at 70 days. The abbreviations are as follows: WT - wild type, VC - vector control, L1 to L5 - transgenic lines L1 to L5.

Following a 10-day recovery period post-salinity stress, transgenic plants demonstrated a rapid recuperative response, as evidenced by the emergence of one to two new leaves per plant. Notably, transgenic plants completed their life cycle within 70 days of the recovery phase, marked by precocious flowering and seed maturation. However, these plants did not attain comparable height, growth, and seed pod production relative to the control condition. A similar pattern was observed in the post-drought recovery of transgenic plants, as they manifested early flowering and seed maturation to complete their life cycle.

### Superior ROS homeostasis in transgenics under stress conditions

Quantitative measurements of hydrogen peroxide (H_2_O_2_) and malondialdehyde (MDA) accumulation, were conducted in control and transgenic plants. Under unstressed conditions, no significant differences in H_2_O_2_ concentration were observed between control and transgenic lines (Figure 3A). However, under salinity and drought stress conditions, a considerable increase in H_2_O_2_ concentration was detected in control plants compared to transgenic lines. Similarly, no statistically significant differences in MDA content were found between control and transgenic plants under unstressed conditions (Figure 3B). In contrast, under abiotic stress conditions, lipid peroxidation significantly increased in control plants compared to transgenic lines and unstressed conditions, while the lipid peroxidation status of transgenic plants remained relatively stable under stress conditions.

**Figure 3:**
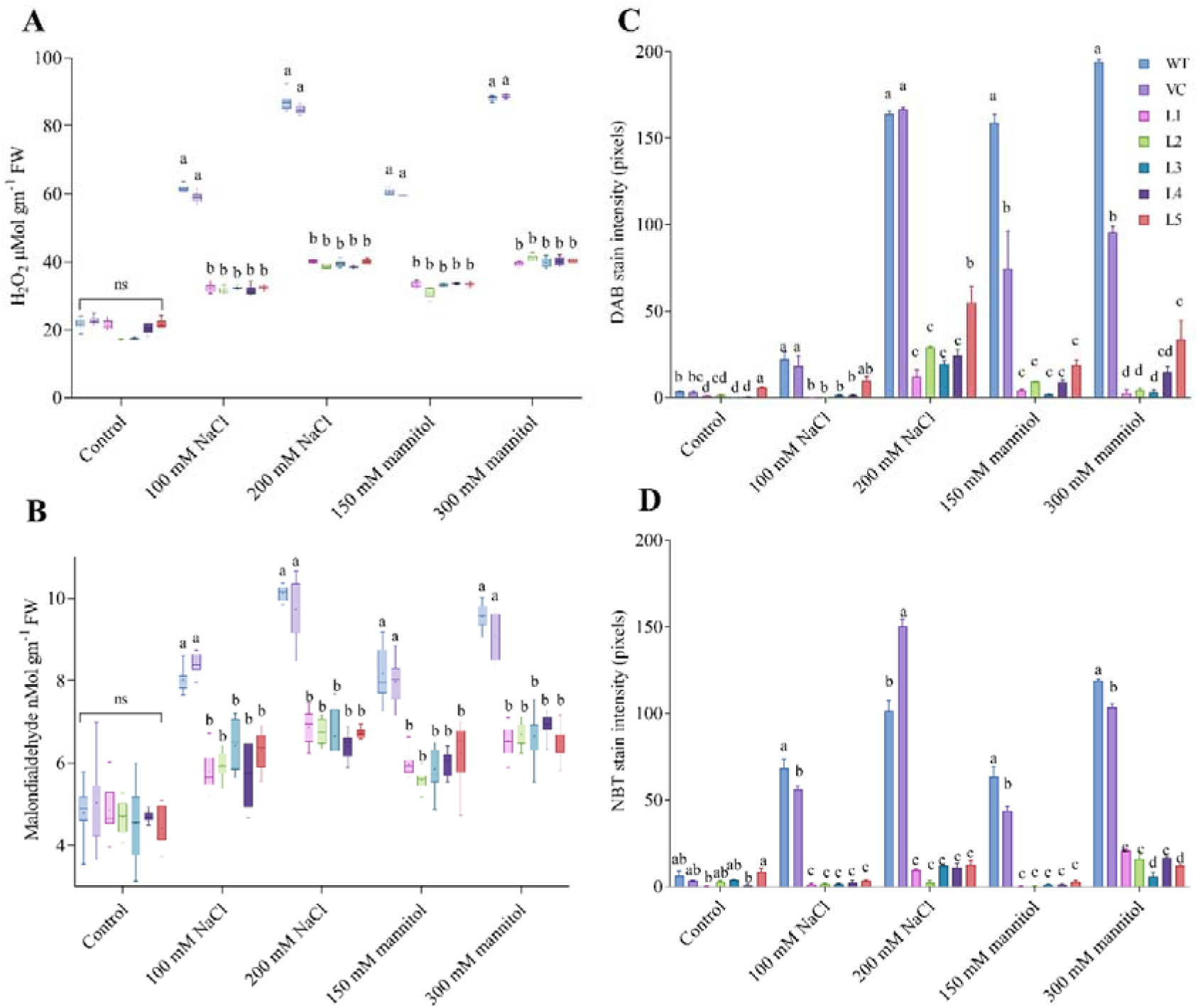
Oxidative stress indicators in plants: This figure consists of: (A) A box plot displaying Hydrogen peroxide concentrations in plants (B) Malondialdehyde concentrations, indicative of lipid peroxidation levels in plants. Data represent means from three independent experiments (n=12). The box plot illustrates the minimum, 25th percentile, median, 75th percentile, and maximum values, with the plus sign indicating the mean value. Stain intensities for: (C) DAB and (D) NBT assays, measured in pixels, assess ROS levels in plants under various abiotic stress conditions. Data are presented as means (±SE) from three independent experiments (n=5). Different lowercase letters denote statistically significant differences at p < 0.05 among different plant lines under the same conditions, determined by ANOVA and Tukey’s HSD test. The term "ns" denotes non-significant differences. Abbreviations include WT - wild type, VC - vector control, and L1 to L5 - transgenic lines L1 to L5.

Histochemical qualitative estimation of ROS, such as H_2_O_2_ and free oxygen radicals, was performed using 3,3’-diaminobenzidine (DAB) and nitroblue tetrazolium (NBT) staining (Figure 4). Under unstressed conditions, all plants, including control and transgenic lines, exhibited faint staining or brown precipitates, indicative of normal physiological levels of ROS. However, under all abiotic stress conditions, control plants displayed significantly higher brown staining of DAB compared to transgenic lines. Maximum accumulation of H2O2, as evidenced by brown precipitates, was observed in control plant samples under 200 mM NaCl and 300 mM mannitol stress conditions. Similar results were obtained with NBT staining for in vivo detection of free oxygen radicals, with control plants showing significantly higher staining by blue precipitates under stress conditions compared to transgenic lines. Quantitative estimation of DAB and NBT stained pixels using ImageJ software yielded results consistent with biochemical measurements, revealing lower intensities of DAB and NBT staining in transgenic plants under abiotic stress conditions, which indicated lower accumulation of H_2_O_2_ and superoxide radicals compared to control plants (Figure 3C-D).

**Figure 4:**
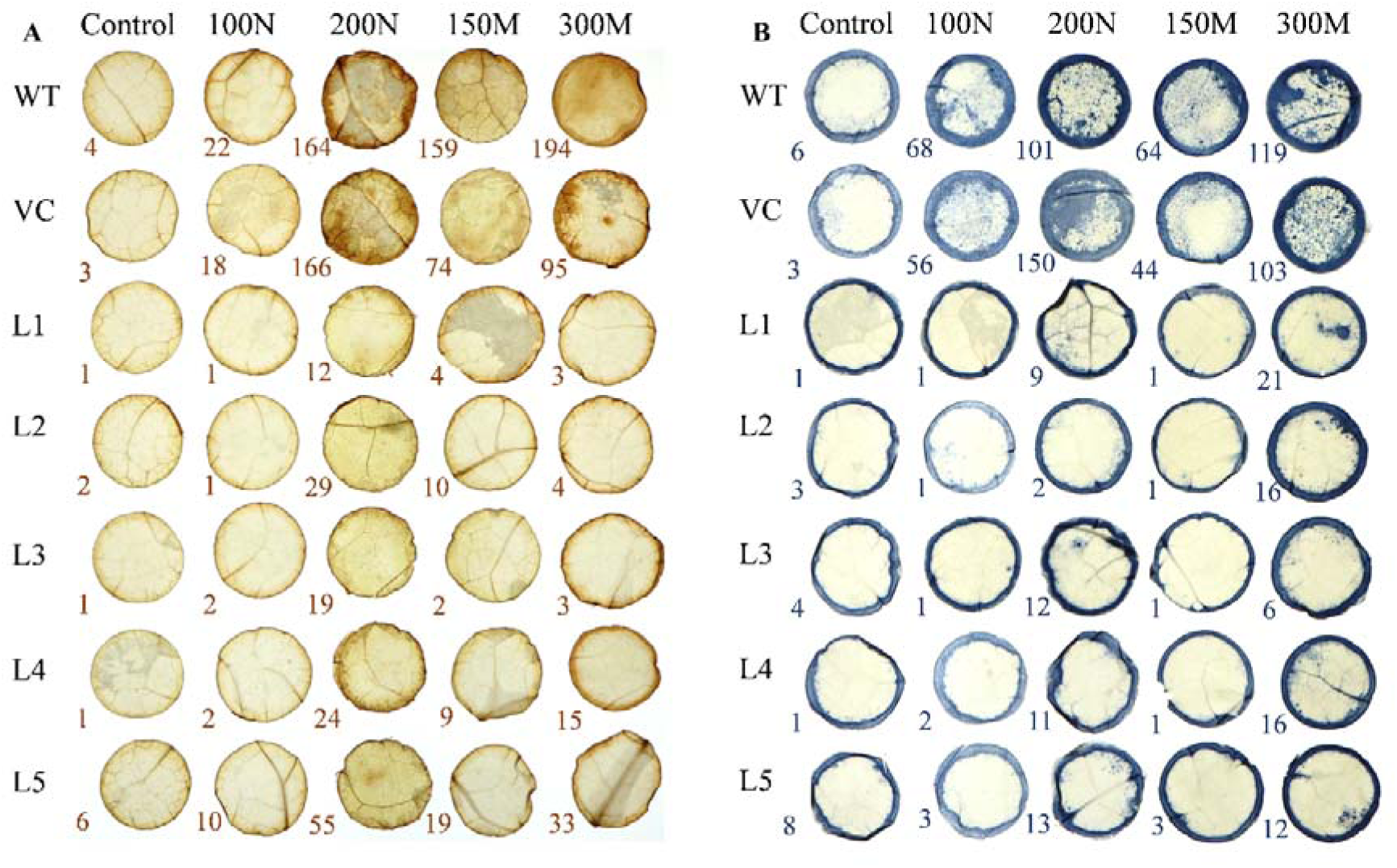
I*n Vivo* detection of reactive oxygen species (ROS) using DAB and NBT assays: This figure demonstrates the detection of ROS in leaf tissue via DAB (3,3’-Diaminobenzidine) and NBT (Nitro Blue Tetrazolium) staining. Leaf discs, 15 mm in diameter, were cut using a sharp cutter and incubated in their respective DAB and NBT staining solutions. To facilitate the visualisation of staining spots, the leaf discs were subsequently bleached with ethanol. The intensity of the staining spots was quantified using ImageJ software, excluding the outer 2 mm diameter to mitigate the impact of tissue damage from cutting. Displayed images are representative of three independent experiments, each with at least five replicates. The abbreviations include: WT - wild type, VC - vector control, L1 to L5 - transgenic lines L1 to L5, with control for unstressed conditions, 100N for 100 mM NaCl stress, 200N for 200 mM NaCl stress, 150M for 150 mM mannitol stress, and 300M for 300 mM mannitol stress. Stain intensity values, measured using ImageJ software, are provided in the lower left corner of each image.

### Molecular mechanism of improving abiotic stress tolerance depicted by transcriptomics and metabolomics

Microarray analysis identified differentially expressed transcripts (≥ 5-fold, p < 0.05) in transgenic plants compared to wildtype plants under unstressed, 200 mM NaCl, and 300 mM mannitol stress conditions. Under unstressed conditions, a total of 1,376 differentially expressed genes (DEGs) were identified in transgenic plants compared to wildtype plants, in which 595 DEGs were upregulated and 780 DEGs were downregulated (Figure S2B). In the case of salinity stress, 2,822 DEGs were identified in transgenic plants, in which with 2,096 DEGs were upregulated and 726 DEGs were downregulated (Figure S2C). Finally, under drought stress conditions, 2,322 DEGs were identified, with 1,394 DEGs were upregulated and 928 DEGs were downregulated (Figure S2D). The Venn diagram illustrates the overlap of differentially expressed genes across control, salinity, and drought stress conditions in transgenic plants (Figure S2A). The diagram quantifies the unique and shared differentially expressed genes between the three conditions. A total of 1,199 genes were exclusively regulated under unstressed conditions, 2,446 under salinity stress, and 1,966 under drought stress. Additionally, 84 genes were shared between unstressed and salinity, 64 between unstressed and drought, and 263 between salinity and drought conditions.

Gene ontology and enrichment analysis revealed the biological processes influenced by the overexpression of the SbPIP2 gene in transgenic plants under both unstressed and abiotic stress conditions. Scatter plots in semantic space (Figure 5A-C) demonstrated that, under unstressed conditions, biological processes such as photosynthesis, seed germination, embryo development, phosphate ion transport, regulation of cell growth, flower development and xylem and phloem formation were significantly impacted. In comparison, under salinity stress, responses to abscisic acid, calcium-mediated signalling, potassium ion transport, ethylene biosynthetic processes, photosynthetic processes, root epidermal cell differentiation, sugar-mediated signalling pathways, and regulation of stomatal movement were markedly affected. Likewise, under drought stress, processes including calcium-mediated signalling, intracellular transport, photosynthesis, gibberellin biosynthesis, responses to osmotic and oxidative stress, reproduction, cytoskeleton organisation, and flower development were most influenced.

**Figure 5:**
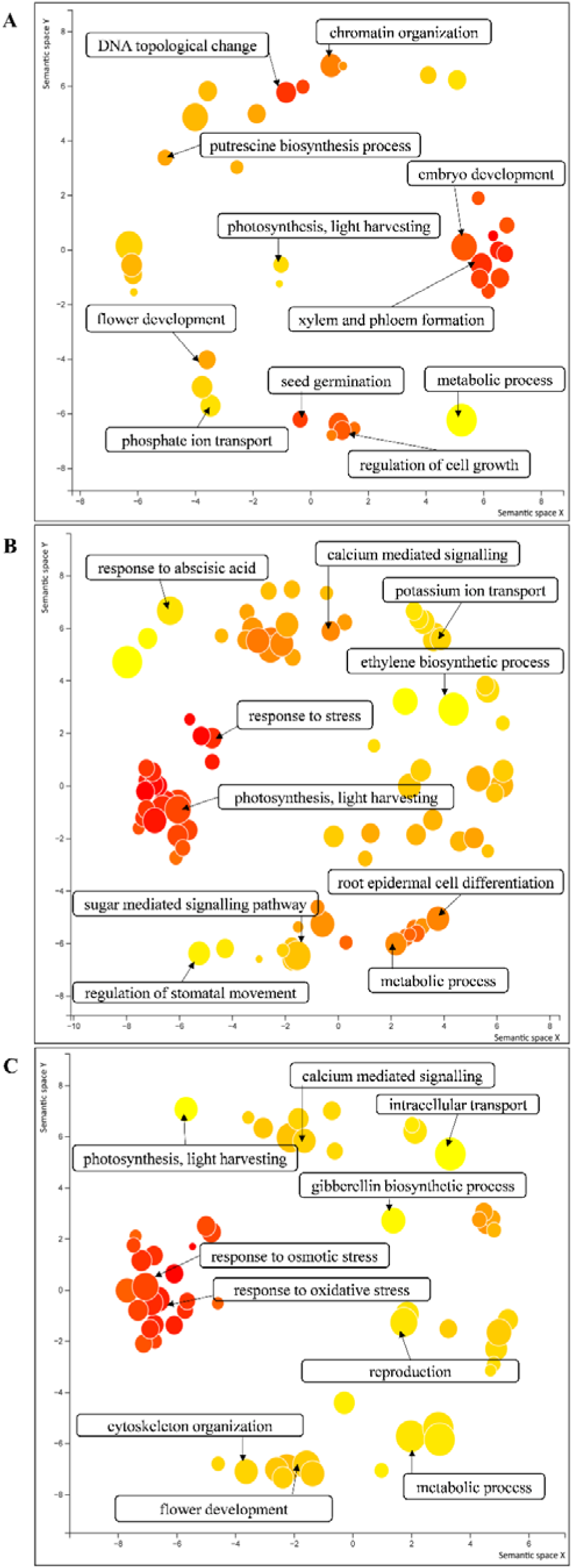
Changes in biological processes of transgenic vs. wildtype plants depicted using gene ontology across different conditions: (A) Under unstressed conditions, (B) Under 200 mM NaCl stress, and (C) Under 300 mM mannitol (simulating drought) stress. Multidimensional scaling was employed to reduce the complexity of the data, plotting values in a semantic space. The size of each circle represents the log fold change in expression, while the color gradient from yellow to orange indicates an abundance of related transcripts.

Important curated DEGs involved in transgenic plant growth, development, and abiotic stress tolerance are listed in Table 1 with relative fold change and respected p values. Genes involved in abscisic acid (ABA) biosynthesis, catabolism and ABA-mediated stomatal closure signalling were differentially expressed under both salinity and drought stress conditions in transgenic plants. Similarly, transcripts associated with ethylene biosynthesis, spermidine biosynthesis, auxin biosynthesis and root development, shikimic acid pathway, and circadian rythym and flowering control were differentially expressed in transgenic plants compared to control plant. Detailed molecular mechanism of improving abiotic stress tolerance depicted by transcriptomics is described in discussion section.

**Table 1:** List of curated genes involved in plant growth, development, and abiotic stress response from transcriptomics data.

| Transcript Cluster ID | Gene name | Gene function | UniProt ID | Unstressed condition |  | Salinity stress (200 mM NaCl) |  | Drought stress (300 mM mannitol) |  |
| --- | --- | --- | --- | --- | --- | --- | --- | --- | --- |
|  |  |  |  | Fold Change (Transgenic vs. WT) | ANOVA p-value | Fold Change (Transgenic vs. WT) | ANOVA p-value | Fold Change (Transgenic vs. WT) | ANOVA p-value |
| Absciscic acid (ABA) biosynthesis and catabolism |  |  |  |  |  |  |  |  |  |
| NtPMIa1g88013e1_st | 9-cis-epoxycarotenoid dioxygenase NCED2, chloroplastic | Conversion of 9-cis-violaxanthin and 9-cis-neoxanthin to ABA biosynthesis precursor xanthoxin | FH950147 | - | - | 7.26 | 0.03 | - | - |
| NtPMIa1g83222e2_st | 9-cis-epoxycarotenoid dioxygenase NCED4 |  | ET970393 | - | - | 5.82 | 0.01 | - | - |
| NtPMIa1g26575e1_st | 9-cis-epoxycarotenoid dioxygenase NCED1 |  | FG164902 | - | - | - | - | 7.01 | 0.04 |
| NtPMIa1g83697e1_st | 9-cis-epoxycarotenoid dioxygenase NCED9, chloroplastic |  | FH945887 | - | - | - | - | 5.58 | 0.03 |
| NtPMIa1g12914e1_st | Absciscic acid 8'-hydroxylase 1 (Cytochrome P450 707A5) | Conversion of ABA to biologically inactive ABA-glucosyl esters | FH480426 | - | - | -7.32 | 0.01 | -8.84 | 0.01 |
| NtPMIa1g132948e1_st | Absciscic acid 8'-hydroxylase 2 (Cytochrome P450 707A6) |  | ET912943 | - | - | -13.45 | 0.01 | -16.69 | 0.04 |
| NtPMIa1g3192e1_st | UDP-glycosyltransferase 71E1 |  | FH213402 | - | - | -16.62 | 0.01 | -7.76 | 0.03 |
| NtPMIa1g4034e1_st | UDP-glycosyltransferase 73E1 |  | FI078848 | - | - | -33.14 | 0.01 | -27.70 | 0.04 |
| NtPMIa1g104803e2_x_st | Absciscic acid 8'-hydroxylase 4 (ABA 8'-hydroxylase 4) (EC 1.14.14.137) (Cytochrome P450 707A4) |  | FH982799 | - | - | - | - | -6.94 | 0.02 |

| **Protein phosphatase 2Cs (PP2Cs) (core signalling module for ABA mediated stomatal closure)** |  |  |  |  |  |  |  |  |  |
| --- | --- | --- | --- | --- | --- | --- | --- | --- | --- |
| NtPMIa1g214<br>805e1_st | *ABA INSENSITIVE 1*<br>(ABI1) | Inhibits OST1<br>(open stomata 1)<br>in the absence of<br>ABA to prevent<br>stomatal closure |  | - | - | -8.73 | 0.04 | - | - |
| NtPMIa1g846<br>39e2_st | *ABA INSENSITIVE 2*<br>(ABI2) | Inhibits GHR1<br>(guard cell<br>hydrogen<br>peroxide resistant<br>1) in the absence<br>of ABA to<br>prevent stomatal<br>closure | FH308017 | - | - | -13.3 | 0.01 | -18.38 | 0.01 |
| NtPMIa1g446<br>64e1_st | *Protein phosphatase 2C 12* | Inhibits SnRKs<br>(SNF1-related<br>protein kinases) in<br>the absence of<br>ABA to prevent<br>stomatal closure | FI055221 | 5.54 | 0.03 | - | - | - | - |
| NtPMIa1g483<br>78e2_s_st | *Protein phosphatase 2C 40* |  | ET973631 | 5.80 | 0.04 | - | - | - | - |
| NtPMIa1g747<br>34e1_st | *Protein phosphatase 2C 43* |  | FG186456 | 11.65 | 0.04 | -6.15 | 0.02 | - | - |
| NtPMIa1g988<br>43e1_st | *Protein phosphatase 2C 48* |  | FH989456 | 11.19 | 0.03 | - | - | 14.44 | 0.01 |
| NtPMIa1g812<br>67e1_s_st | *Protein phosphatase 2C 52* |  | FH349151 | 10.79 | 0.01 | - | - | - | - |
| NtPMIa1g200<br>466e1_s_st | *Protein phosphatase 2C 63* |  | EH622268 | - | - | -10.51 | 0.04 | -7.20 | 0.03 |
| NtPMIa1g103<br>550e1_s_st | *Protein phosphatase 2C 78* |  | FH945363 | - | - | -42.84 | 0.01 | -39.86 | 0.04 |
| NtPMIa1g814<br>47e2_s_st | *Protein phosphatase 2C 37* |  | ET869561 | - | - | -16.47 | 0.01 | -10.00 | 0.04 |
| **Ethylene biosynthesis** |  |  |  |  |  |  |  |  |  |
| NtPMIa1g181<br>173e1_s_st | *S-adenosylmethionine*<br>*synthase* | Catalysed L-Methionine to S-Adenosylmethioni ne (SAM) | FG148581 | - | - | 9.29 | 0.02 | 10.93 | 0.01 |
| NtPMIa1g64585e1_st | *ACC synthase* | Catalysed SAM to 1-aminocyclopropane-1-carboxylate (ACC) | FH430844 | - | - | -12.10 | 0.04 | -9.24 | 0.03 |
| NtPMIa1g180753e1_st | *ACC oxidase 1* | Catalysed ACC to ethylene | EH617872 | - | - | - | - | -54.46 | 0.04 |
| NtPMIa1g47710e2_st | *ACC oxidase 2* |  | FG200599 | - | - | -23.06 | 0.03 | - | - |
| NtPMIa1g48907e1_s_st | *ACC oxidase 3* |  | FG193081 | - | - | -32.06 | 0.04 | - | - |
| NtPMIa1g190129e2_x_st | *AP2-like ethylene-responsive transcription factor AIL6* | *Ethylene-responsive transcription factors* | ET042791 | - | - | -6.02 | 0.02 | -8.21 | 0.04 |
| NtPMIa1g32023e2_s_st | *Ethylene-responsive transcription factor ABR1-like* |  | FH048685 | - | - | -16.85 | 0.02 | - | - |
| NtPMIa1g61179e1_st | *Ethylene response factor 4* | Inhibits ethylene biosynthesis | FH962950 | - | - | 7.43 | 0.03 | 7.9 | 0.01 |
| **Spermidine biosynthesis** |  |  |  |  |  |  |  |  |  |
| NtPMIa1g142930e1_s_st | *S-adenosylmethionine decarboxylase 3* | Conversion of SAM to decarboxy-S-adenosylmethionine (dcSAM) | FH463775 | - | - | 7.38 | 0.02 | 8.18 | 0.01 |
| NtPMIa1g53560e2_st | *Spermidine synthase (SPDSY), Polyamine aminopropyltransferase (PAPT)* | Conversion of dcSAM to spermine and spermidine | FH111569 | 5.03 | 0.02 | 27.14 | 0.01 | 25.77 | 0.01 |
| NtPMIa1g149920e1_st | *Polyamine oxidase 3* | Interconversion of spermine spermidine and putrescine |  | - | - | 5.98 | 0.01 | 9.7 | 0.02 |
| NtPMIa1g141<br>164e1_st | *Thermospermine synthase* | Synthesis of thermospermine | FH232500 | 6.18 | 0.01 | 5.48 | 0.01 | 6.02 | 0.02 |
| Auxin biosynthesis and root development |  |  |  |  |  |  |  |  |  |
| NtPMIa1g172<br>461e1_st | *Enoyl-CoA delta isomerase 2, peroxisomal-like* | *Involved with IBR1 and IBR3 in the peroxisomal beta-oxidation of indole-3-butyric acid (IBA) to form indole-3-acetic acid (IAA), a biologically active auxin* | FG183308 | 22.14 | 0.03 | - | - | - | - |
| NtPMIa1g184<br>416e1_st | Auxin conjugate hydrolase (IAA-amino acid hydrolase ILR1-like protein) | function in storage, inactivation or inhibition of the growth regulator auxin | FH708980 | - | - | -6.43 | 0.02 | - | - |
| NtPMIa1g196<br>139e1_st | *WALLS ARE THIN1 (WAT1),* | plant-specific protein that dictates secondary cell wall thickness of wood fibres | ET924591 | 242.75 | 0.03 | 153.07 | 0.01 | 78.94 | 0.03 |
| NtPMIa1g107<br>139e1_st | *WAT1-related protein* |  | FI022414 | 114.66 | 0.01 | 109.86 | 0.03 | 43.28 | 0.02 |
| NtPMIa1g142<br>852e1_st | *Auxin-responsive GH3-like protein 5 (Indole-3-acetic acid-amido synthetase GH3.5)* | linked to lateral root development | FH530447 | - | - | 7.20 | 0.03 | - | - |
| NtPMIa1g948<br>51e1_st | *Indole-3-acetic acid-amido synthetase GH3.11* |  | FH241629 | - | - | - | - | 7.87 | 0.03 |
| NtPMIa1g121<br>109e1_st | *Indole-3-acetic acid-amido synthetase GH3.10* |  | FI069924 | - | - | - | - | 7.04 | 0.01 |
| Shikimic acid pathway |  |  |  |  |  |  |  |  |  |
| NtPMIa1g71645e5_st | *Prephenate dehydratase* | Production of aromatic amino acids and folates | FG197758 | - | - | - | - | 6.30 | 0.01 |
| NtPMIa1g187686e1_st | *Chorismate mutase 3, chloroplastic* |  | FH736002 | - | - | 7.92 | 0.02 | 8.52 | 0.01 |
| NtPMIa1g215183e1_st | *DAHPS, 3-deoxy-D-arabino-2-heptulosonate 7-phosphate synthase* |  | FG197758 | 9.32 | 0.01 | 5.91 | 0.01 | 12.62 | 0.01 |
| Circadian rhythm and flowering control |  |  |  |  |  |  |  |  |  |
| NtPMIa1g19982e1_st | Flowering time control protein FCA | Plays a major role in the promotion of the transition of the vegetative meristem to reproductive development | FH621309 | -15.29 | 0.00 | - | - | - | - |
| NtPMIa1g100647e1_st | *PSEUDO-RESPONSE REGULATOR 5 (PRR5)* | part of the repressor system that governs the biological clock in plants | FH274108 | 66.54 | 0.04 | 8.20 | 0.03 | 78.04 | 0.01 |
| NtPMIa1g40744e1_st | *LNK1-like protein* | Functions as a transcriptional coactivator of PRR5 | ET760143 | 49.68 | 0.03 | 6.04 | 0.01 | 10.31 | 0.01 |
| NtPMIa1g75901e1_st | FRIGIDA-like protein | Circadian rhythm controlling protein | FG140151 | -5.58 | 0.03 | - | - | - | - |
| NtPMIa1g92446e1_st | *TIME FOR COFFEE* |  | FH035513 | - | - | -5.90 | 0.01 | -5.33 | 0.01 |

Metabolomic profiling of transgenic and wildtype (WT) plants under unstressed and abiotic stress conditions was conducted using GC-MS. Metabolite concentrations were determined using Ribitol as an internal standard and analysed using MetaboAnalyst version 5.0 (https://www.metaboanalyst.ca/). This analysis identified a total of 108 unique metabolites across all samples (Table 2). Venn diagram analysis was employed to visualise the differential accumulation of metabolites under control and stress conditions (Figure S3A-B). This revealed a consistent presence of 21 to 25 metabolites in all samples. Specifically, under salinity and drought stress, transgenic plants uniquely accumulated 12 and 19 metabolites, respectively. Conversely, 11 metabolites under salinity stress and 10 under drought stress were common to both WT and transgenic plants, but absent under unstressed conditions.

**Table 2:**
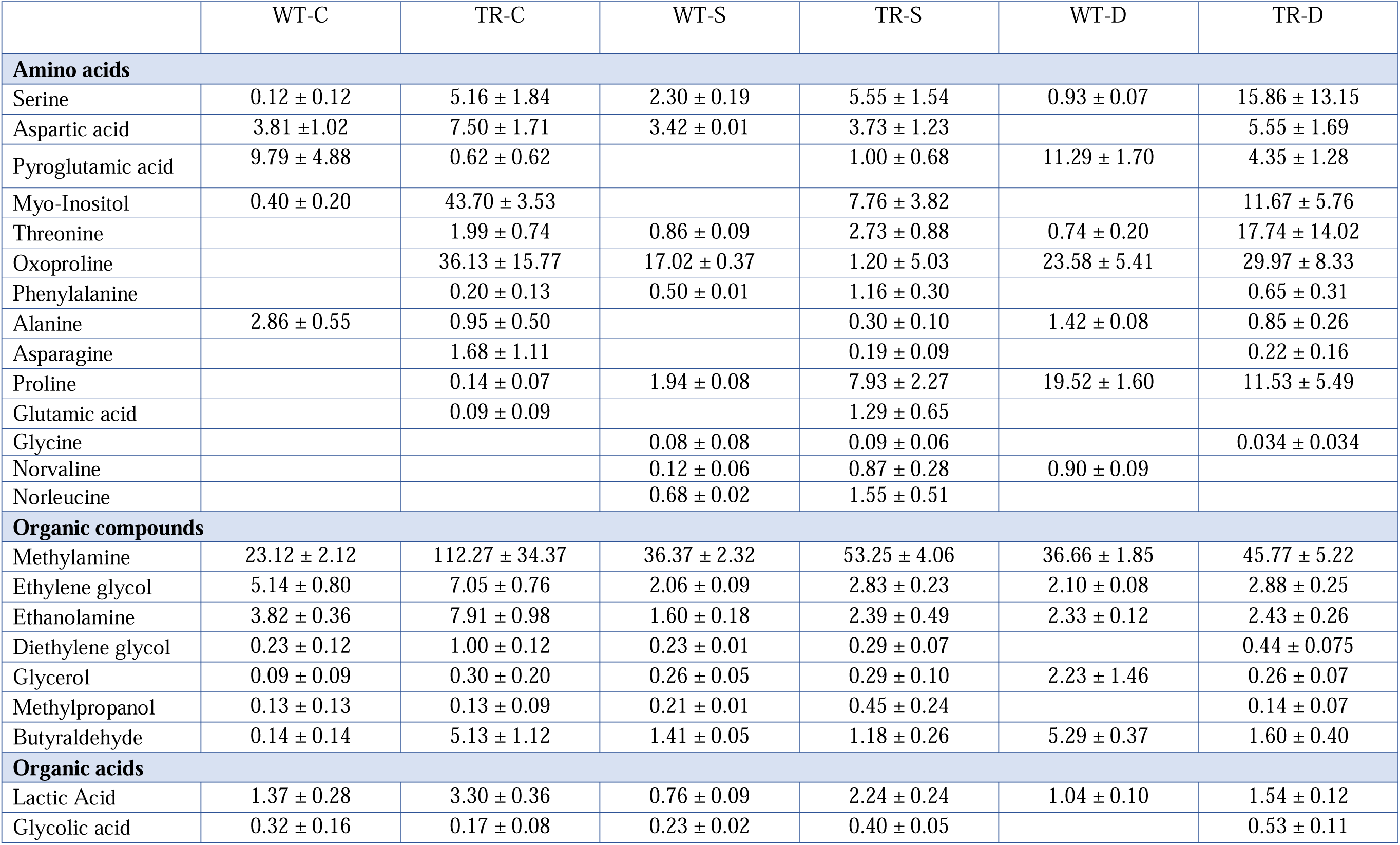
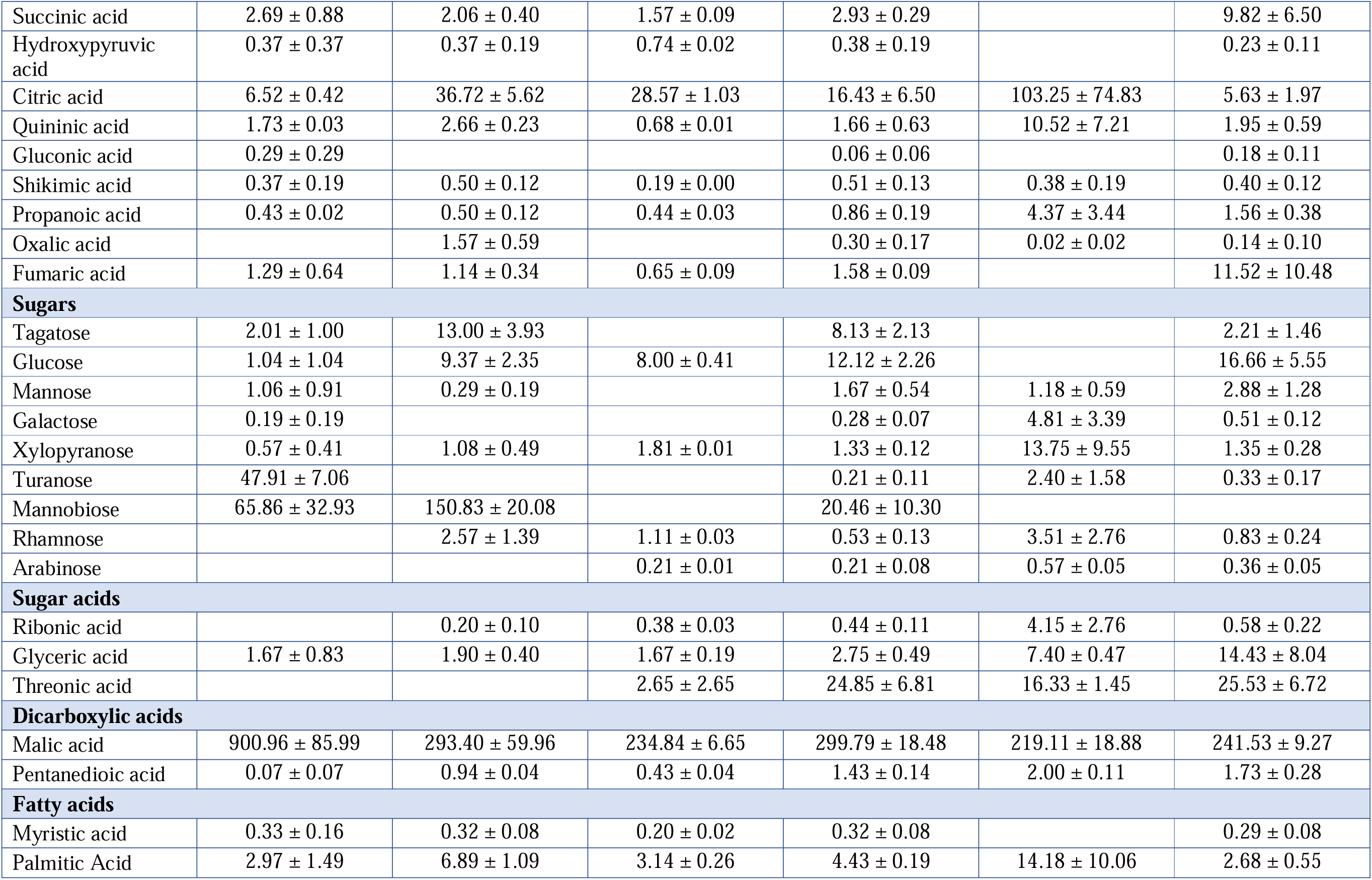
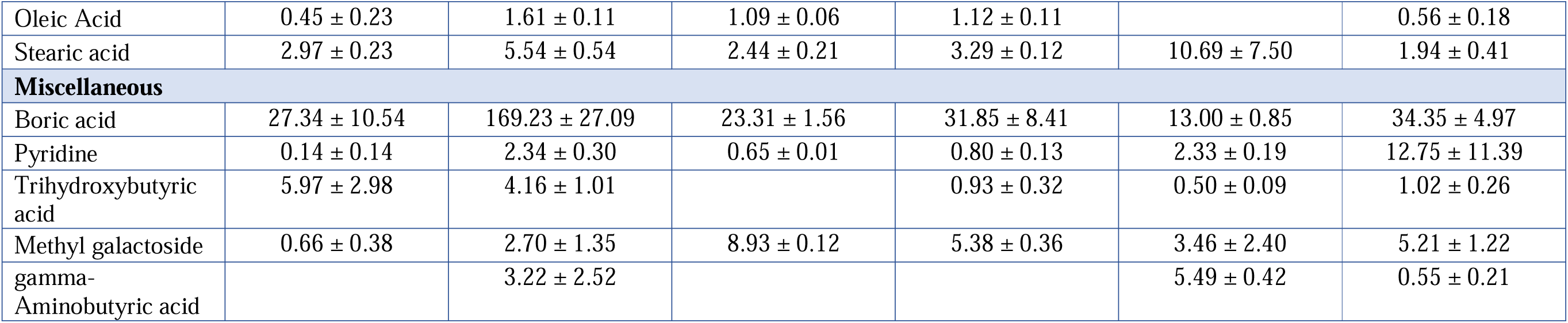
List of selected metabolites with their concentration identified by untargeted metabolomics.

Relative changes in metabolite concentrations were quantified using a log 2-fold change threshold via MetabolAnalyst. Under control conditions, transgenic plants showed an increase in 8 metabolites, a decrease in 5, and 39 metabolites remained unchanged compared to WT plants (Figure S3C). Under salinity stress, 7 metabolites decreased, while 49 showed no significant change (Figure S3D). During drought stress, 8 metabolites significantly increased, 19 decreased, and 27 remained unchanged in transgenic plants relative to WT (Figure S3E). The heatmap highlighted significant shifts in metabolite concentrations in transgenic plants under salinity and drought stress (Figure 6). Under salinity stress, increases were observed in norleucine, glutamic acid, methylpropanol, phenylalanine, and glucose, while decreases were noted in gluconic acid, myristic acid, boric acid, and mannobiose. Under drought stress, increases in fumaric acid, succinic acid, glucose, threonine, and serine were detected, alongside decreases in aspartic acid, glycine, norleucine, hydroxy pyruvic acid, and glutamic acid.

**Figure 6:**
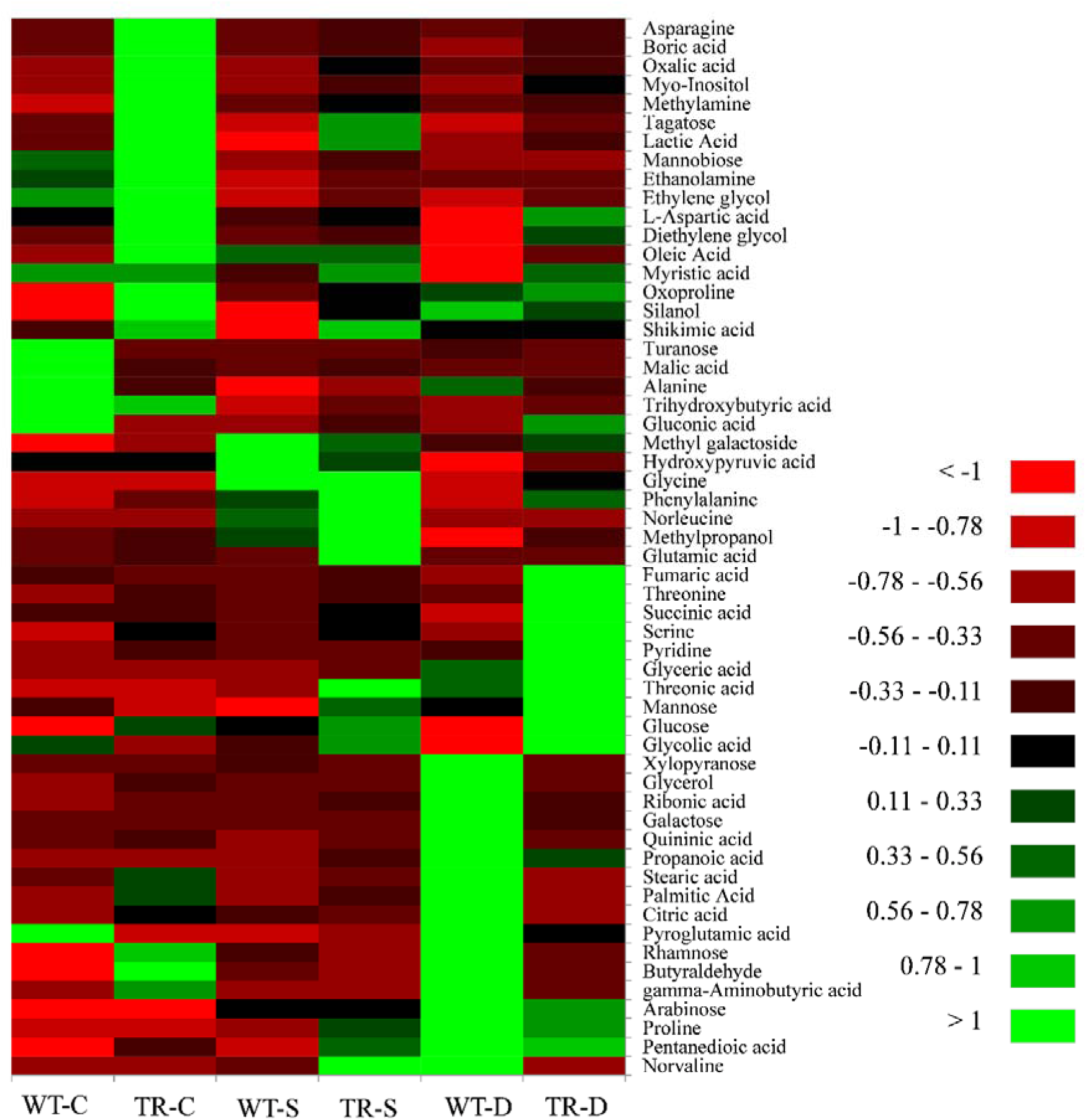
Heatmap of metabolite concentrations from untargeted metabolomics: This heatmap presents the concentration of selected metabolites identified through untargeted metabolomics. For improved visualisation, metabolite concentrations were normalised to a scale from 1 to -1. The labels are as follows: WT-C denotes wildtype plants under unstressed conditions, TR-C represents transgenic plants under unstressed conditions, WT-S for wildtype plants under 200 mM NaCl stress, TR-S for transgenic plants under 200 mM NaCl stress, WT-D for wildtype plants under 300 mM mannitol (drought) stress, and TR-D for transgenic plants under 300 mM mannitol (drought) stress.

Partial Least Squares Discriminant Analysis (PLS-DA) indicated a higher overall accumulation of metabolites such as glycolic acid, palmitic acid, lactic acid, diethylene glycol, ethylene glycol, threonine, gluconic acid, shikimic acid, and malic acid (Figure 7).

**Figure 7:**
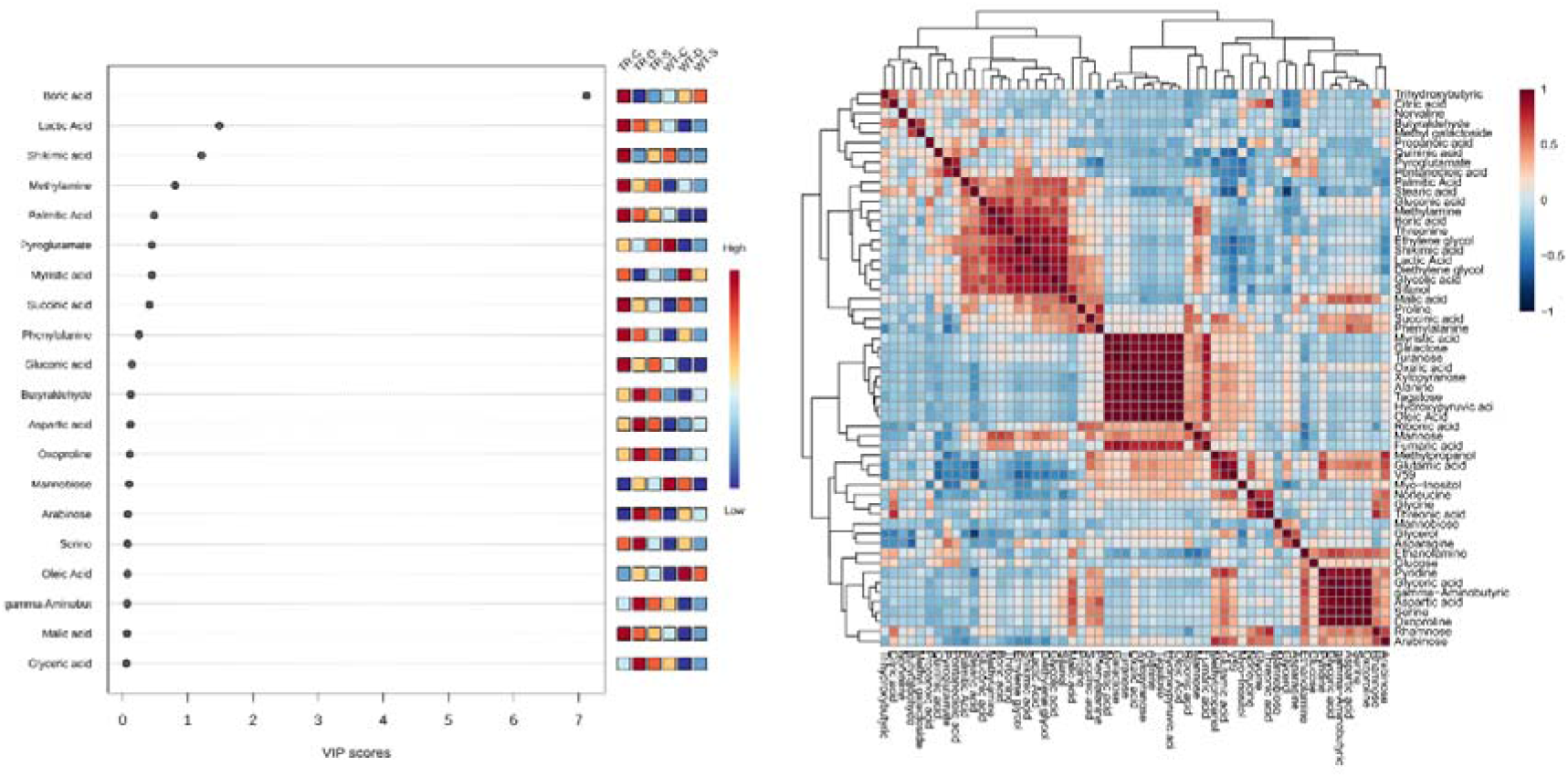
Partial least squares discriminant analysis (PLS-DA) and correlation map of metabolites: This figure showcases a partial least squares discriminant analysis and a correlation map for selected metabolites identified through untargeted metabolomics. The conditions are labeled as follows: WT-C for wildtype plants under unstressed conditions, TR-C for transgenic plants under unstressed conditions, WT-S for wildtype plants under 200 mM NaCl stress, TR-S for transgenic plants under 200 mM NaCl stress, WT-D for wildtype plants under 300 mM mannitol (drought) stress, and TR-D for transgenic plants under 300 mM mannitol (drought) stress.

Alpha hydroxy acids like glycolic and lactic acid were more abundant in both transgenic and control plants under unstressed conditions, but reduced under stress. In contrast, aspartic acid and myo-inositol accumulated more under stress conditions. Glyceric acid levels rose in transgenic plants but fell in controls under stress. The correlation map showed positive associations between certain monosaccharides and TCA cycle intermediates, and among specific amino acids (Figure 7). Conversely, correlations were observed between certain fatty acids and metabolites like shikimic acid, glycolic acid, gluconic acid, and lactic acid.

Pathway topology analysis revealed that under unstressed conditions, the glyoxylate and dicarboxylate metabolism pathways were most impacted (Figure S4). Other significantly affected pathways included aminoacyl-tRNA biosynthesis, amino acid biosynthesis, unsaturated amino acid biosynthesis, cyanoamino metabolism, butanoate metabolism, and pathways for cutin, suberine, and wax biosynthesis. Under abiotic stress conditions, aminoacyl-RNA biosynthesis was the most affected pathway. Under salinity stress, glyoxylate and dicarboxylate metabolism also showed significant impact, along with amino acid metabolism, galactose metabolism, cyanoamino metabolism, unsaturated fatty acids biosynthesis, and glutathione metabolism. Similarly, under drought stress, the most impacted pathways were galactose metabolism and others, including amino acid metabolism, cyanoamino metabolism, ascorbate and aldarate metabolism, unsaturated fatty acids biosynthesis, and pentose and glucuronate interconversions.

## Discussion

Plants must tightly regulate their water status throughout their lifecycle, especially during growth and seed germination, under varying environmental conditions like abiotic stresses. This regulation is mainly achieved through stomatal control and water movement at cellular and tissue levels (Maurel et al., 2021). Aquaporins, present in plant cell membranes, are key in cell-to-cell water transport and osmotic regulation (Patel and Mishra, 2021). Here we used combined qualitative and quantitative approach ranging from histochemical staining to transcriptomics, to gain a detailed understanding of the processes that enable plants to tolerate environmental stresses by overexpression of *SbPIP2* gene. This discussion aims to integrate these findings, providing a clear view of the underlying mechanisms through which genetic modifications in transgenic plants lead to improved resistance to challenging environmental conditions.

Plants maintain delicate balance of ROS in cell by generation as by-product of metabolism and scavenging activities. ROS play important role in plant abiotic stress as signalling molecule (Gilroy et al., 2016). Qualitative histochemical ROS staining by DAB and NBT; and quantitative H_2_O_2_ and lipid peroxidation estimation showed differential accumulation of ROS and H_2_O_2_ under control and transgenic plants. Transgenic plants showed significantly less accumulation of ROS compared to control (WT and VC) plants under abiotic stress treatments. Similarly, *Arabidopsis* plants ectopically expressing *ThPIP2;5* and *MsPIP2;2* showed significantly less accumulation of ROS compared to WT plants under abiotic stress treatments (Li et al., 2019; Wang et al., 2018) Aquaporins play an important role in complex metabolism of seed germination process. Seed germination process starts with uptake of water by dry seeds in which water channel like aquaporins plays a major role. In rice silencing of *OsPIP1;3* and *OsPIP1;1* genes reduced whereas overexpressing same genes increased germination rate of rice seedling indicating relation of aquaporins and seed germination parameters (Liu et al., 2007). Similar results were found in our study where under unstressed condition, seed germination parameters were similar in control (WT and VC) and transgenic plants but germination percentage and germination energy were significantly higher and relative stress injury was significantly lower in transgenic plants compared to control (WT and VC) under all abiotic stress conditions. Microarray results also confirmed that seed germination, embryo development, regulation of cell growth and other metabolic process involved in seed germination and viability were affected in transgenic plants compared to control plants. Similarly, transgenic *Arabidopsis* expressing *JcPIP2;7* and *JcTIP1;3* showed higher seed germination rate, improved seed yield and seed viability compared to WT plants under salinity and drought stress (Khan et al., 2015).

Though hydroponics culture system has better repeatability, plants have different growth pattern in soil compared to hydroponics culture condition. Transgenic plants expressing *SbPIP2* showed significantly faster growth, early flowering and seed settling compared to control (WT and VC) plants in soil. Similar results were found in transgenic *Arabidopsis* overexpressing *ZxPIP1;3* gene showing higher growth of transgenics under control and salinity stress condition (Li et al., 2021). After 25 days of salinity and drought stress transgenic plants were less effected by stress compared to control plants. Similar results were found in transgenic potato lines expressing *StPIP1* gene under water deficit conditions (Wu et al., 2009).

After stress treatments, recovery also play a major role in plants survival, completion of lifecycle and yield. After 10 days of recovery, transgenic plants started sprouting new leaves. Long term recovery of 70 days after abiotic stresses showed significantly improvement in the physiology of transgenic plants. In fact, transgenic plants showed flowering, seed settling and completion of life cycle. Similarly, transgenic banana plants overexpressing *MusaPIP1;2* gene showed faster recovery after drought stress compared to WT plants (Sreedharan et al., 2015). Transgenic *Arabidopsis* expressing *TsPIP1;2* also showed 26% higher survival rate and post stress recovery compared to WT plants (Li et al., 2018). Additionally, microarray results showed that transcript related to response to oxidative stress, calcium and sugar mediated signaling pathway, regulation of stomatal movement, phytohormone biosynthesis, photosynthesis, flower development and reproduction were differentially expressed in transgenic plants under abiotic stresses compared to control plants. Similarly, untargeted metabolomics and pathway topology analysis supported differential accumulation of metabolites related to amino acid and sugar metabolism.

Transcriptomics analysis of control and transgenic plants under unstressed and stressed condition reveal mechanism by which transgenic plants enhanced their abiotic stress resilience. Plant hormone ABA known to regulates growth and development especially under a/biotic stress conditions. Two homologues of ABA biosynthesis gene *9-cis-epoxycarotenoid dioxygenase, NCED2* and *NCED4* were significantly upregulated in salinity stress whereas NCED1 and NCED9 were significantly upregulated in drought stress in transgenic plants. In contrast, ABA catabolising genes like *Abscisic acid 8’-hydroxylase 1* and *UDP-glycosyltransferase 71E1* were decreased significantly in transgenic plants under both abiotic stresses. This indicates increased ABA accumulation by increased biosynthesis and decreased catabolism in transgenic plants compared to wildtype plants under abiotic stresses. Similar results were found in transgenic wheat where overexpression of TaHsfA6f protein leads to increased ABA level and enhanced abiotic stress resilience (Bi et al., 2020). In the absence of ABA, ABI1 (ABA INSENSITIVE 1) and ABI2 (ABA INSENSITIVE 2) interacts with OST1/GHR1 and inhibit stomatal closure process (Hua et al., 2012). Expression of ABI1 and ABI2 was reduced in transgenic plants under both abiotic stress condition. Protein phosphatase 2C (PP2C) group proteins interacts with SnRKs (SNF1-related protein kinases) leading to inhibition of stomatal closure (Chen et al., 2021). Transgenic plants showed reduced expression of PP2Cs like PP2C37, PP2C43, PP2C63, and PP2C78 under salinity and drought stress conditions. These results suggest that under abiotic stress stomatal aperture were more reduced in transgenic plants compared to wildtype plants for reduced water loss and stress survival. In contrast, under unstressed condition PP2C12, PP2C40, PP2C43, PP2C48 and PP2C52 were overexpressed in transgenic plants indicating wider stomatal aperture in transgenic plants for improved stomatal conductance and photosynthesis.

Expression of *S-adenosylmethionine synthase* gene catalysing methionine to SAM (S-Adenosylmethonine) was significantly increased in transgenic plants under both abiotic stress conditions. This serve as a first step in ethylene biosynthesis pathway (Pattyn et al., 2021). Though, expression of two other important enzymes *ACC (1-aminocyclopropane-1-carboxylate) synthase* and *ACC oxidase* were significantly reduced in transgenic plants under stress conditions. Further expression of the *Ethylene response factor 4*, which inhibits ethylene production by epigenetic regulations also increased in transgenic plants. *Ethylene responsive transcription factors ABR1-like* and *AP2-like* were also decreased in transgenic plants under abiotic stresses. These results showed that ethylene concentration was decreased in transgenic plants under abiotic stresses. Previously it was reported that ethylene inhibits ABA mediated stomatal closure pathway (Song et al., 2023). However, here in transgenic plant reduced ethylene promote ABA mediated stomatal closure and reduced water loss under abiotic stress conditions. Also accumulated SAM in the first step of ethylene biosynthesis was diverted to spermidine biosynthesis.

Polyamines, including putrescine, spermidine, and spermine, boost plant growth and stress tolerance, whether applied externally or produced endogenously in genetically modified plants (Tyagi, et al., 2023). The expression of *S-adenosylmethionine decarboxylase 3* (*SAMDC3*), which converts SAM to decarboxy-S-adenosylmethionine (dcSAM), showed a notable increase in transgenic plants subjected to salinity and drought stress conditions. *Spermidine synthase* (*SPDS*), responsible for the biosynthesis of spermidine and spermine, exhibited elevated expression levels in transgenic plants even under baseline conditions, and further increased under abiotic stress compared to both control and wildtype plants. Additionally, the activities of *Polyamine oxidase 3* and *Thermospermine synthase*, key enzymes in polyamine interconversion and thermospermine synthesis (Takano et al., 2012), respectively, were significantly enhanced in transgenic plants under salinity and drought stress relative to their wildtype counterparts. These findings suggest that the transgenic plants demonstrate an upregulated production of polyamines such as putrescine, spermidine, spermine, and thermospermine, contributing to improved tolerance against abiotic stress factors.

ABA dependent activation of the IAA signaling pathway under abiotic stress conditions was shown by ABA-deficient mutants (Zhang et al., 2022). Efficient production, transport, and storage of IAA, vital for plant growth, are influenced by peroxisomal Enoyl-CoA delta Isomerase enzymes through the β-oxidation of IBA into free IAA (Zolman et al., 2008, Li et al., 2019). In the transgenic plants studied, expression levels of the *Enoyl-CoA delta Isomerase 2* (*ECI2*) gene, associated with the biosynthesis of the active form of auxin (indole-3-acetic acid or IAA), were elevated under non-stress conditions. However, these levels remained stable under various abiotic stresses. In the context of salinity stress, there was a noticeable down-regulation of the transcript for *Auxin conjugate hydrolase*, a gene implicated in the deactivation and sequestration of auxin. This observation supports the notion of elevated auxin levels facilitating root growth in transgenic plants as compared to their wildtype counterparts under saline conditions (data not shown). Previous studies in *Arabidopsis* showed that GH3 protein act as a negative regulator in lateral root development and promoting primary root growth (Wang et al., 2023). *Auxin-responsive GH3-like protein 5* under salinity stress, and both *Indole-3-acetic acid-amido synthetase GH3.10* and *GH3.11* under drought conditions were increased in transgenic plants supporting significantly higher primary root length in transgenic plant seedling (data not shown).

Shikimic acid pathway produces chorismate, a precursor for aromatic amino acid production (Tzin and Galili, 2010). Further aromatic amino acid like phenylalanine participates in downstream phenypropanoid pathway to synthesise important antioxidant flavonoids (Kumar et al., 2023). A distinct pattern of gene expression related to the shikimic acid pathway was noted in transgenic plants compared to wildtype plants. Specifically, the transcript levels of *3-deoxy-D-arabino-2-heptulosonate 7-phosphate synthase* (*DAHPS*), a key enzyme in the initial stages of shikimic acid biosynthesis, showed a marked increase in transgenic plants under both control and abiotic stress conditions. Furthermore, *Chorismate mutase 3*, involved in the conversion of chorismate to prephenate, displayed elevated expression levels during salinity and drought stress conditions in transgenic variants. Additionally, there was a considerable rise in the expression of *prephenate dehydratase*, catalysing the formation of phenylalanine from prephenate, particularly under drought stress. Moreover, untargeted metabolomics confirmed that phenylalanine concentration was significantly higher in transgenic plants compared to control plants under salinity and drought stress. These observations collectively indicate an enhanced capability for aromatic amino acid biosynthesis in transgenic plants when subjected to abiotic stress, compared to their wildtype counterparts.

As shown previously transgenic plants showed significant faster growth rate and early flowering compared to wild type plants. Transcriptomics revealed differential expression of major gene involved in circadian rhythm and flowering control. Expression of Flowering time control gene *flowering locus C* (*FLC*) which act as a repressor of flowering in plants was significantly reduced in transgenic plants. FRIGIDA-like protein is a nuclear protein which upregulate FLC and delay flowering in plants. (Chao et al., 2013) This gene was also found downregulated in transgenic plants under control condition can link lower FLC expression and delayed flowering. *Pseudo response regulator 5* (*PRR5*) gene was found to be controlling the circadian rhythm of plants and involved in early flowering of plants. (Fujimori et., 2005) Transcript of *PRR5* was also found increased significantly in transgenic plants under control and unstressed conditions. Also, transcript of LNK1 like protein which act as a transcriptional coactivator of *PRR5* was increased significantly in transgenic plants under control and stressed conditions. *Arabidopsis* mutants showing lower expression of *TIME FOR COFFEE* (*tic*) gene which is a circadian regulator showed increased drought tolerance. (Sanchez[Villarreal et al., 2013). Similarly here in transgenic plants under both salinity and drought abiotic stress *tic* gene was reduced significantly.

In conclusion, our research presents a subtle view of transgenic plant abiotic stress tolerance mechanisms, integrating the reduced ROS levels, enhanced seed germination, and improved plant physiology in both hydroponic and soil conditions due to overexpression of *SbPIP2*. Our comprehensive study has shed light on the multifaceted nature of plant abiotic stress tolerance mechanisms. Through transcriptomics, we have elucidated the pathways through which transgenic plants enhance their resilience against abiotic stresses, predominantly focusing on the ABA, ethylene, and polyamine pathways alongside auxin signalling and the shikimic acid pathway. These findings underscore the importance of hormonal regulation and metabolic adjustments in stress response and reveal the underlying genetic modifications that contribute to enhanced stress tolerance. The altered expression of key genes involved in circadian rhythm and flowering control further demonstrates the extensive influence of these genetic modifications. Recently, Arabidopsis PIP1 and PIP2 aquaporins were found to regulate ABA mediated stress response via phosphorylation and interactor proteins like SnRK2s (Grondin et al., 2015; Rodrigues et al., 2017). However, it is the question of studying how *SbPIP2* interact with ABA signalling networks and modifies transcript networks to increase growth rate and abiotic stress resilience. Overall, our research provides a deeper understanding of the molecular mechanisms underpinning plant abiotic stress tolerance, offering valuable insights for future efforts to improve crop resilience in the face of escalating environmental challenges.

## Acknowledgement

Authors JP, NKG, and BC duly acknowledge Junior Research Fellowship from UGC, New Delhi. Author JP duly acknowledges funding for Newton Bhabha PhD placement program from DBT, New Delhi and British Council, UK. We extend our gratitude to Dr. Chaitanya Joshi from the Gujarat Biotechnology Research Centre for his valuable assistance with the microarray experiment. We also thank Kaksha Savliya and Neha for their help in microarray data annotation.

## Supplimentary Figures

**Figure S1:**
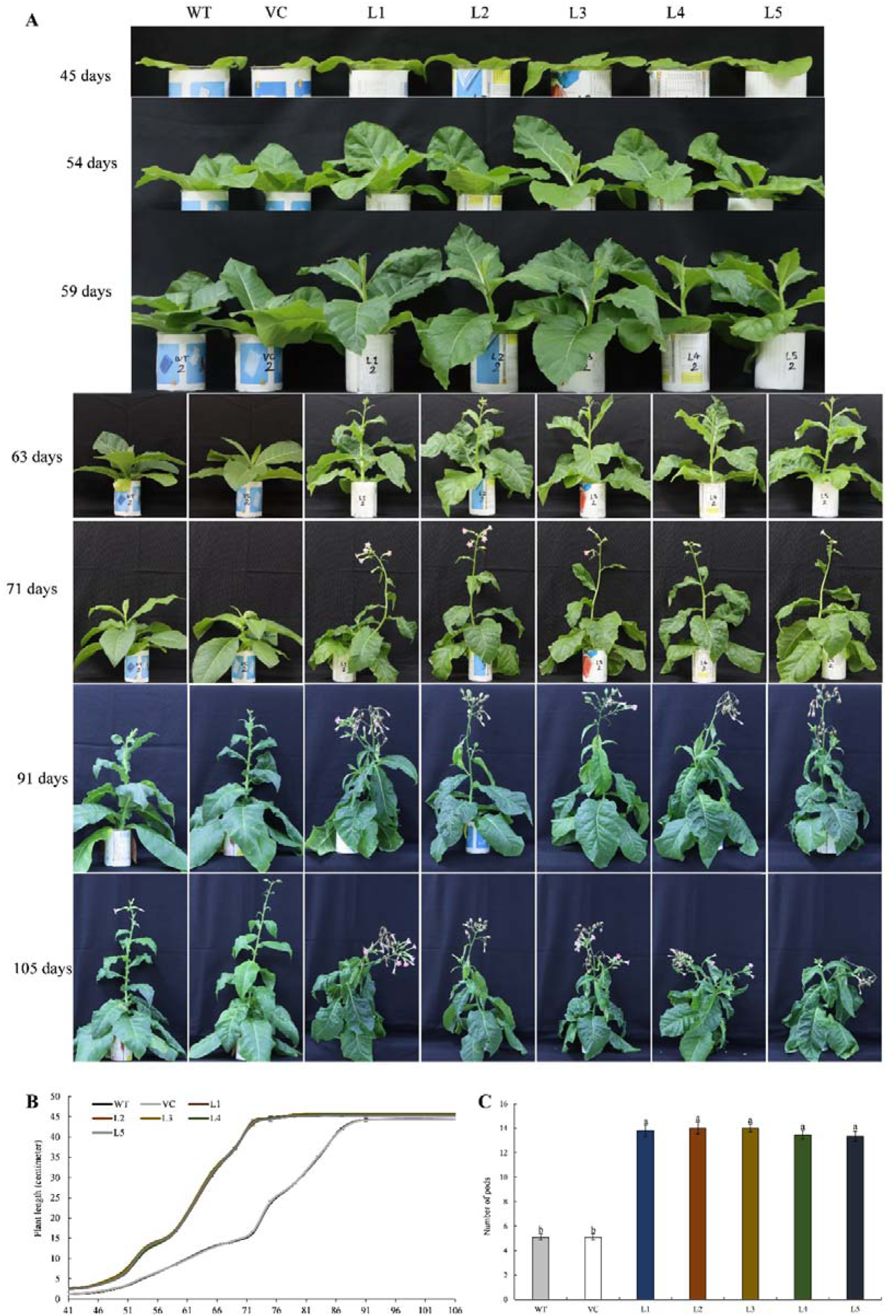
Growth and development analysis of control and transgenic plants (A) This panel shows a growth analysis of control and transgenic plants under hydroponic conditions, documented at regular intervals to highlight differences in growth rates under unstressed conditions. The age of the plants, starting from seed germination, is indicated in days on the left side of the figure. (B) Displays the change in plant length (in centimeters) measured every 4 days. (C) Presents the number of seed pods harbored by 120-day-old mature plants. Values are expressed as means ± SE (n=5). Different lowercase letters indicate statistical differences at p < 0.05 among plant lines, as determined by Tukey’s HSD test.

**Figure S2:**
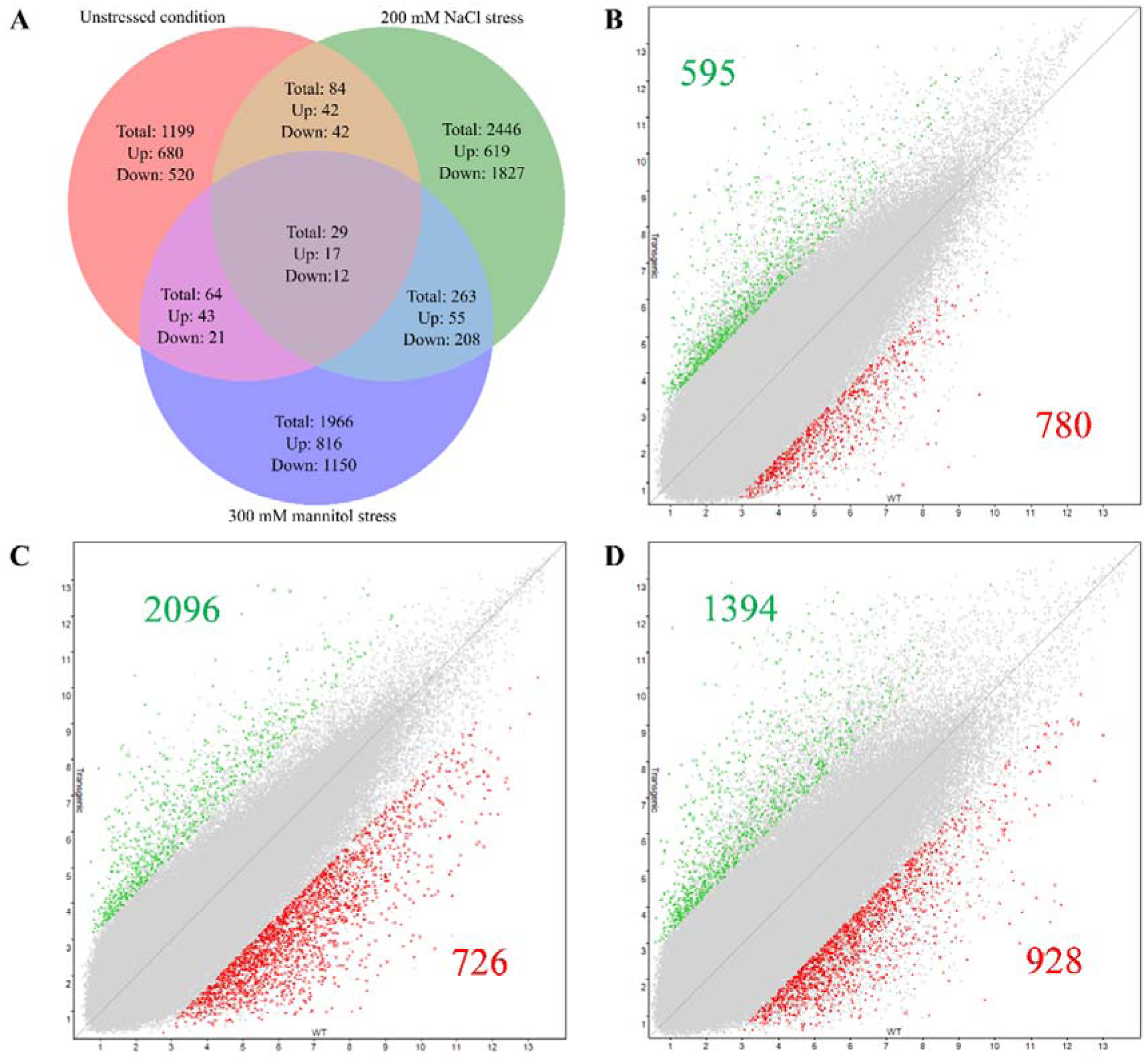
Differential gene expression analysis. (A) A Venn diagram showing the overlap of differentially expressed genes under unstressed, salinity, and drought stress conditions in transgenic plants compared to wildtype plants. (B-D) Scatter plots of differentially expressed genes in transgenic plants compared to wildtype plants under unstressed conditions (B), 200 mM NaCl stress (C), and 300 mM mannitol stress (D), with a minimum threshold of log 5-fold change. Green numbers denote upregulated genes, and red numbers indicate downregulated genes.

**Figure S3:**
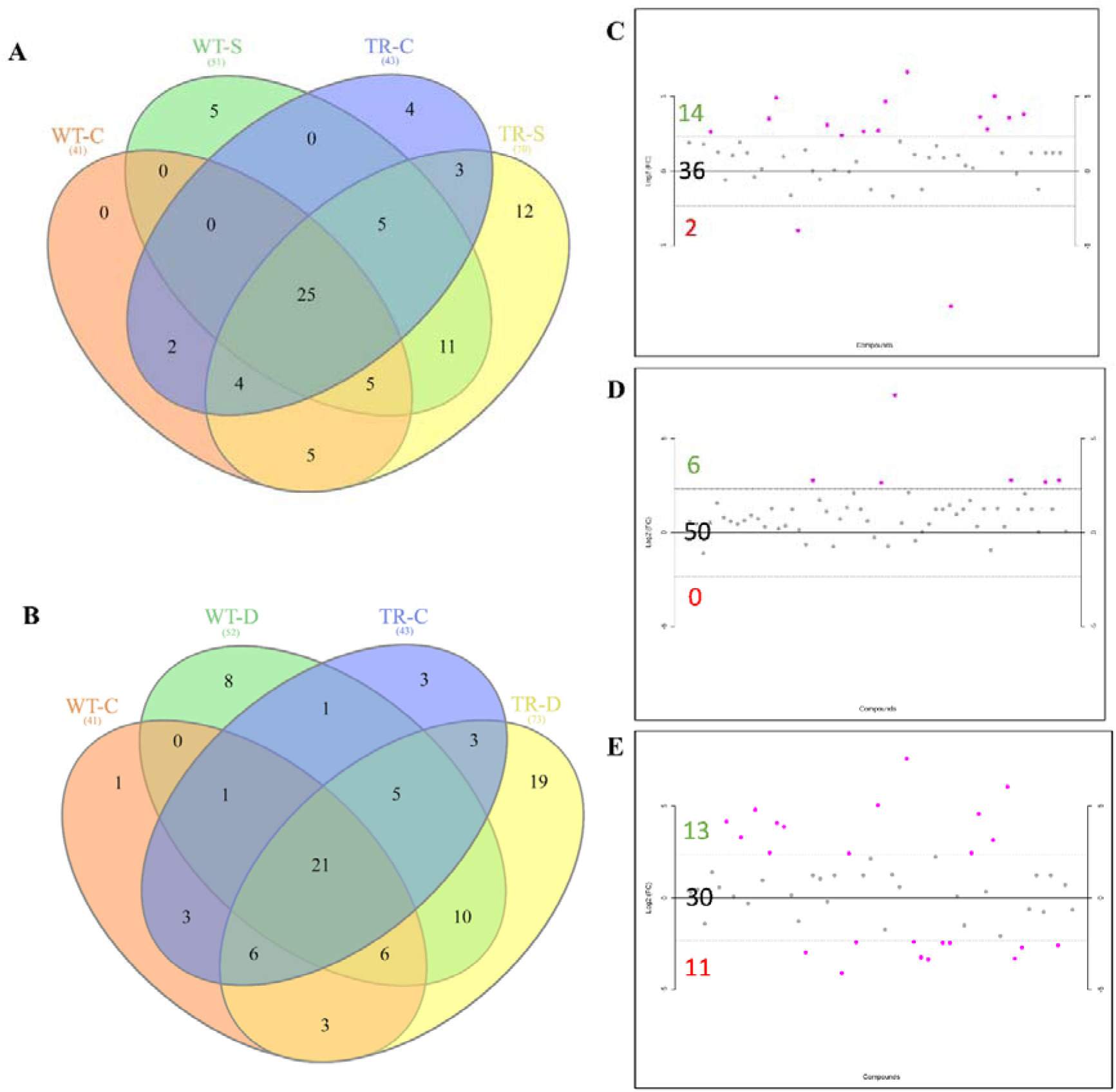
Metabolite accumulation analysis: (A-B) Venn diagrams illustrating the overlap of differentially accumulated metabolites under 200 mM NaCl stress and 300 mM mannitol stress. Labels include WT-C for wildtype in control condition, WT-S for wildtype under 200 mM NaCl stress, TR-C for transgenic plants in unstressed condition, and TR-S for transgenic plants under 200 mM NaCl stress, WT-D for wildtype under 300 mM mannitol stress, and TR-D for transgenic plants under 300 mM mannitol stress. (C-E) Scatter plots depicting differentially accumulated metabolites under unstressed condition €, 200 mM NaCl stress (D), and 300 mM mannitol stress €. Green letters indicate an increase in metabolites, black letters for no significant change, and red letters for a decrease, with a minimum threshold of log 2-fold change.

**Figure S4:**
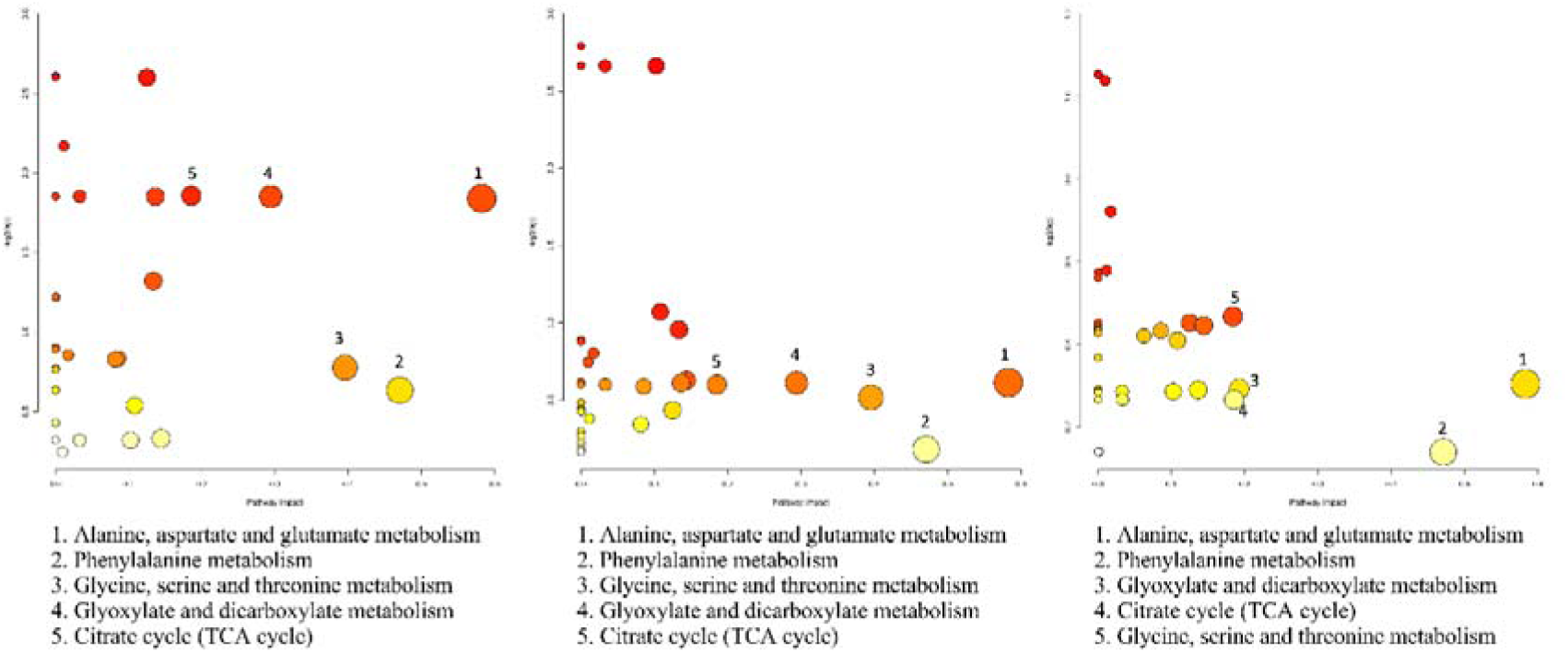
Pathway topology analysis via MetaboAnalyst: This figure presents pathway topology analyses performed using MetaboAnalyst software under unstressed conditions (A), 200 mM NaCl stress (B), and 300 mM mannitol stress (C). The top five metabolism pathways identified are listed below their respective plots.

## Notes

### Competing Interest Statement

The authors have declared no competing interest.

